# Human iPSC-Derived Cardiomyocytes are Susceptible to SARS-CoV-2 Infection

**DOI:** 10.1101/2020.04.21.051912

**Authors:** Arun Sharma, Gustavo Garcia, Vaithilingaraja Arumugaswami, Clive N. Svendsen

**Affiliations:** Board of Governors Regenerative Medicine Institute, Cedars-Sinai Medical Center, Los Angeles, CA 90048, USA; Smidt Heart Institute, Cedars-Sinai Medical Center, Los Angeles, CA 90048, USA; Department of Molecular and Medical Pharmacology, David Geffen School of Medicine, University of California, Los Angeles, Los Angeles, CA 90095, USA; Eli and Edythe Broad Center of Regenerative Medicine and Stem Cell Research, University of California, Los Angeles, Los Angeles, CA 90095, USA

**Author notes:** **Corresponding authors and email addresses:**. **Lead Contact:** Clive N. Svendsen, PhD., Address: 127 S. San Vicente Blvd., Pavilion, Room 8405, Los Angeles, CA 90048.

**Keywords:** COVID-19, SARS-CoV-2, coronavirus, induced pluripotent stem cells, viral myocarditis, cardiomyocytes, cardiovascular biology, heart, stem cell, cardiology

## Abstract

Coronavirus disease 2019 (COVID-19) is a viral pandemic caused by the severe acute respiratory syndrome coronavirus 2 (SARS-CoV-2). COVID-19 is predominantly defined by respiratory symptoms, but cardiac complications including arrhythmias, heart failure, and viral myocarditis are also prevalent. Although the systemic ischemic and inflammatory responses caused by COVID-19 can detrimentally affect cardiac function, the direct impact of SARS-CoV-2 infection on human cardiomyocytes is not well-understood. We used human induced pluripotent stem cell-derived cardiomyocytes (hiPSC-CMs) as a model system to examine the mechanisms of cardiomyocyte-specific infection by SARS-CoV-2. Microscopy and immunofluorescence demonstrated that SARS-CoV-2 can enter and replicate within hiPSC-CMs, localizing at perinuclear locations within the cytoplasm. Viral cytopathic effect induced hiPSC-CM apoptosis and cessation of beating after 72 hours of infection. These studies show that SARS-CoV-2 can infect hiPSC-CMs *in vitro*, establishing a model for elucidating the mechanisms of infection and potentially a cardiac-specific antiviral drug screening platform.

## INTRODUCTION

Coronavirus disease 2019 (COVID-19), which is caused by the severe acute respiratory syndrome coronavirus 2 (SARS-CoV-2), has been declared an international pandemic, causing the hospitalization and deaths of hundreds of thousands of people worldwide (Ramaiah and Arumugaswami, 2020). SARS-CoV-2, a single-stranded enveloped RNA virus, is known to use the ACE2 receptor to enter host lung tissue, followed by rapid viral replication (Hoffmann et al., 2020). Clinical presentation demonstrates predominantly pulmonary symptoms, including cough, shortness of breath, pneumonia, and acute respiratory distress syndrome (Madjid et al., 2020). However, there is mounting evidence that SARS-CoV-2 infection may cause cardiac complications including elevated cardiac stress biomarkers, arrhythmias, and heart failure (Madjid et al., 2020). A recent study demonstrated significantly elevated troponin levels among some COVID-19 patients, indicating cardiac injury, and notably, cardiac injury was associated with increased risk of mortality (Shi et al., 2020a). The etiology of cardiac injury in COVID-19, however, remains unclear. Cardiac injury may be ischemia-mediated, and the profound inflammatory and hemodynamic impacts seen in COVID-19 have been hypothesized to cause atherosclerotic plaque rupture or oxygen supply-demand mismatch resulting in ischemia (Madjid et al., 2020). Alternatively, cardiac tissue expresses the ACE2 receptor, further suggesting feasibility of direct viral internalization in cardiomyocytes (Chen et al., 2020). Indeed, case studies have raised suspicion for cardiac injury mediated by direct myocardial viral infection and resulting fulminant myocarditis (Hu et al., 2020; Tavazzi et al., 2020; Zeng et al., 2020). In order to gain further insights into the cardiac pathophysiology of COVID-19, it will be critical to determine whether SARS-CoV-2 can directly infect isolated human cardiomyocytes *in vitro*. Elucidating the pathogenic mechanism of cardiac injury in COVID-19 could ultimately guide therapeutic strategies. Antiviral agents could potentially mitigate cardiac complications if the underlying mechanism of cardiac injury is direct myocardial viral infection.

Primary human cardiomyocytes are difficult to obtain and maintain for research use. Improved methods to convert human induced pluripotent stem cells (hiPSCs) to multiple somatic lineages have enabled *in vitro* mass production of patient-specific cells, including hiPSC-derived cardiomyocytes (hiPSC-CMs) (Sharma et al., 2020). The hiPSC-CMs express relevant proteins found in adult human CMs, can spontaneously contract, can be made in weeks using defined differentiation protocols, and can be genetically customized using genome editing (Sharma et al., 2018). HiPSC-CMs express *ACE2*, which increases over 90 days of differentiation (Churko et al., 2018). The hiPSC-CMs are also responsive to inotropic drugs such as norepinephrine, and beating rates can be controlled via electrical stimulation (Toepfer et al., 2019). Because hiPSC-CMs can be purified and replated for downstream applications, research groups in academia and industry now utilize these cells for cardiovascular disease modeling and high-throughput drug screening assays. HiPSC-CMs can recapitulate cellular phenotypes for cardiovascular diseases including various forms of cardiomyopathy (Lan et al., 2013; Sun et al., 2012) and drug-induced cardiotoxicity (Burridge et al., 2016; Sharma et al., 2017). Notably, hiPSC-CMs have also shown promise as an *in vitro* model for studying the mechanisms of direct cardiomyocyte viral infection in the context of viral myocarditis. A previous study demonstrated that coxsackievirus B3 (CVB3), one of the major causative agents for viral myocarditis, can rapidly infect and proliferate within hiPSC-CMs (Sharma et al., 2014). CVB3, like SARS-CoV-2, is a positive-sense, single-stranded RNA virus, although unlike SARS-CoV-2, it does not have a viral envelope. The hiPSC-CMs produce the coxsackie and adenovirus receptor protein, which is needed for infection by CVB3. Detrimental virus-induced cytopathic effect was observed in hiPSC-CMs within hours of CVB3 infection, manifesting in cell death and contractility irregularities. Importantly, this study also established hiPSC-CMs as a cardiac-specific antiviral drug screening platform, and demonstrated that drugs such as interferon beta and ribavirin can stymie CVB3 proliferation *in vitro* (Sharma et al., 2014). Interferon beta was able to transcriptionally activate viral clearance gene networks in CVB3-infected hiPSC-CMs.

Here, the aforementioned foundational viral myocarditis study is extended to assess the effect of SARS-CoV-2 on hiPSC-CMs. We show that hiPSC-CMs are susceptible to SARS-CoV-2 infection, resulting in functional alterations and cytopathic effects. These cellular phenotypes occurred in the absence of inflammatory and hemodynamic impacts. The results provide new mechanistic insights into the cardiac manifestations of COVID-19 and suggest that the heart may be susceptible to direct infection by SARS-CoV-2.

## RESULTS

### Human induced pluripotent stem cells can differentiate into cardiomyocytes

An hiPSC control line (02iCTR) was generated by the Cedars-Sinai Medical Center iPSC Core from peripheral blood mononuclear cells and shown to be fully pluripotent (Laperle et al., 2020). The hiPSCs were differentiated into hiPSC-CMs using an established monolayer differentiation protocol utilizing small molecule modulators of Wnt signaling (Sharma et al., 2015). Differentiated hiPSC-CMs were metabolically purified by depriving cells of glucose, as previously published (Sharma et al., 2015). Purified hiPSC-CMs expressed standard cardiac sarcomeric markers cardiac troponin T (cTnT) and α-actinin **(Figure 1A)**.

**Figure 1:**
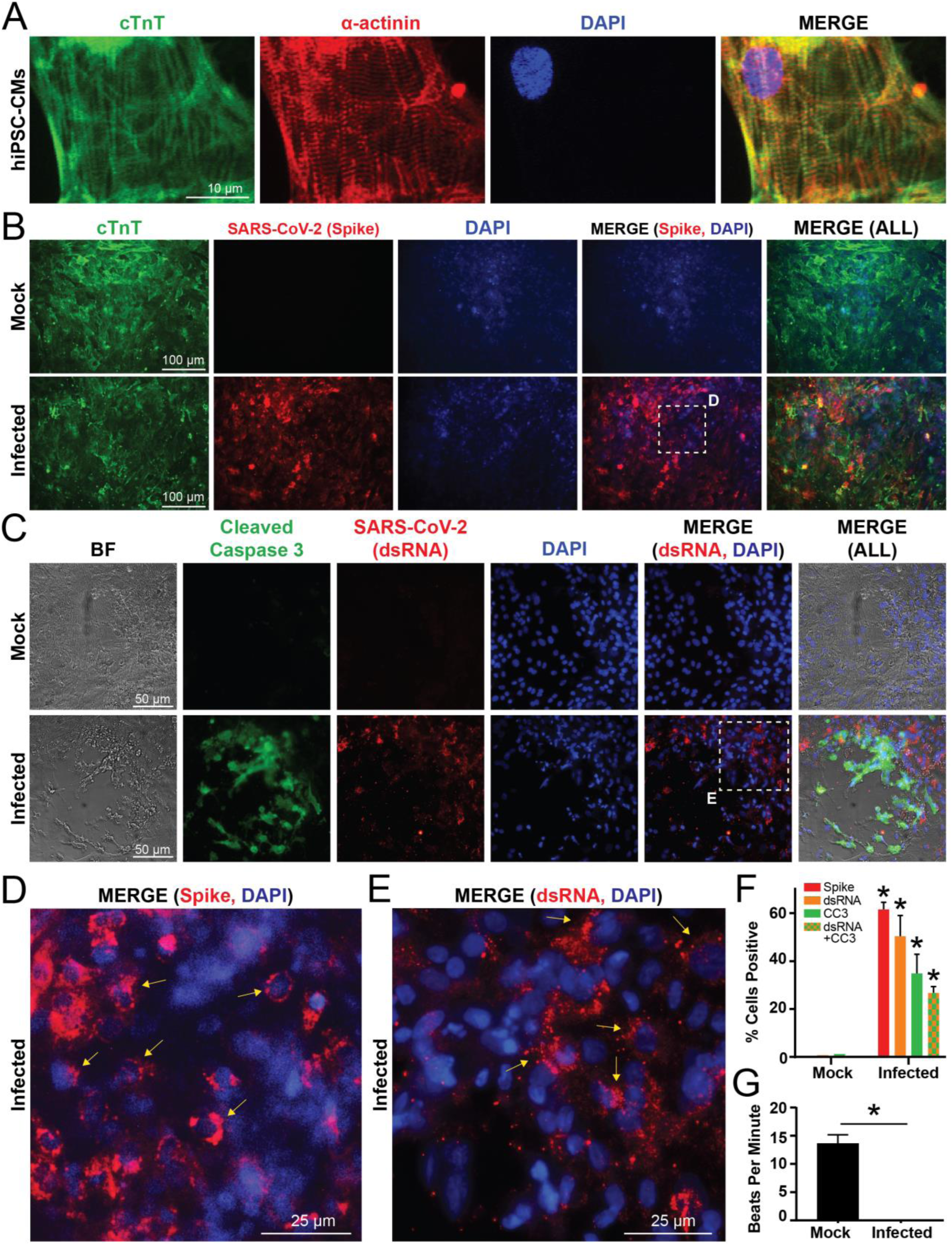
SARS-CoV-2 internalizes and replicates within hiPSC-CMs *in vitro*, eliciting cytopathic effect and contractility alterations. **A)** Human iPSC-CMs exhibit standard sarcomeric markers including cardiac troponin T (cTnT) and α-actinin with DAPI as nuclear counterstain. **B)** Immunofluorescence for cTnT and SARS-CoV-2 “spike” protein demonstrates that hiPSC-CMs can be infected by SARS-CoV-2. SARS-CoV-2 spike protein is not present in mock infected cultures. **C)** HiPSC-CMs after SARS-CoV-2 infection, but not mock infection, exhibit signs of cellular apoptosis, indicated by morphological changes seen in bright field (BF) and cleaved caspase-3 production. A second SARS-CoV-2 antibody marks a viral-specific double-stranded intermediate RNA (dsRNA). **D)** Magnified inset from panel B shows a merged immunofluorescence image for SARS-CoV-2 spike protein and DAPI. Arrows indicate perinuclear accumulation of viral particles, and suggests active viral protein translation and genome replication at perinuclear ribosomes and membranous compartments. **E)** Magnified inset from panel C shows a merged immunofluorescence image for SARS-CoV-2 dsRNA and DAPI. Arrows indicate perinuclear viral replication sites. **F)** Quantification of immunofluorescence indicates percentage of total DAPI-positive cells that are positive for spike protein, viral dsRNA, cleaved caspase-3 (CC3), and dsRNA+CC3 in hiPSC-CMs infected with SARS-CoV-2, compared to mock infection. N=5-7 images quantified for each stain for mock and infected conditions. **G)** Quantification of beats per minute in wells containing hiPSC-CMs with mock infection versus wells containing hiPSC-CMs infected with SARS-CoV-2. N=6 videos recorded for each condition. See Supplemental Movie 1 for representative video clips. * indicates p < 0.05.

### Purified hiPSC-CMs can be directly infected by SARS-CoV-2

Purified hiPSC-CMs were replated into 96-well plates at 100,000 cells per well and allowed to regain contractility **(Supplemental Movie 1)** before being subjected to SARS-CoV-2 infection. The SARS-CoV-2 was obtained from the Biodefense and Emerging Infections (BEI) Resources of the National Institute of Allergy and Infectious Diseases (NIAID) and titered on Vero-E6 cells (see Methods). The hiPSC-CMs were infected with SARS-CoV-2 at a multiplicity of infection (MOI) of 0.1 for 72 hours, with a mock treatment without virus serving as a control condition. In order to determine if the virus could enter and proliferate within hiPSC-CMs, cells from both infected and mock conditions were stained for cardiac marker cardiac troponin T (cTnT) and SARS-CoV-2 viral capsid “spike” protein **(Figure 1B)**. The infected hiPSC-CMs stained positively for spike protein, suggesting that SARS-CoV-2 can establish active infection in hiPSC-CMs.

### SARS-CoV-2 infection of hiPSC-CMs causes apoptosis and cessation of beating

To determine if SARS-CoV-2 induced a cytopathic effect on hiPSC-CMs, mock and infected hiPSC-CMs were stained for the apoptosis marker cleaved caspase-3, as well as for the double stranded RNA (dsRNA) intermediate unique to positive sense RNA virus infection **(Figure 1C)**. The dsRNA and spike protein stains represent two independent assays for visualizing SARS-CoV-2 viral uptake and genome replication in hiPSC-CMs. A proportion of infected cells were positive for dsRNA and also stained positive for cleaved caspase-3, indicating that hiPSC-CMs were undergoing virus-induced apoptosis. Notably, both the SARS-CoV-2 spike protein and dsRNA intermediates localized at perinuclear regions in hiPSC-CMs **(Figure 1D, 1E)**, consistent with prior results with coronavirus infection in non-cardiomyocytes (Hagemeijer et al., 2012) and coxsackievirus infection on hiPSC-CMs (Sharma et al., 2014). This suggests the presence of a viroplasm whereby hiPSC-CM ribosomal machinery and other membranous components are being co-opted for viral replication and protein translation. Quantification of stained cells demonstrated the percent of total cells that were positive for dsRNA and spike protein, as well as for cleaved caspase-3 **(Figure 1F).** Simple beat rate contractility analysis was also conducted, whereby 30 second videos were taken of wells containing mock and infected hiPSC-CMs **(Supplemental Movie 1)**. Functionally, infected hiPSC-CMs ceased beating after 72 hours of SARS-CoV-2 infection, whereas mock wells continued to contract **(Figure 1G)**. Taken together, these results indicate that hiPSC-CMs are susceptible to SARS-CoV-2 infection and downstream detrimental cytopathic effects, that the SARS-CoV-2 may be able to replicate in distinct perinuclear locations within hiPSC-CMs by co-opting cellular organelles for viral protein translation, and that SARS-CoV-2 infection significantly reduces functional contractility in hiPSC-CMs.

## DISCUSSION

Results presented here establish for the first time that human iPSC-derived cardiomyocytes are susceptible to direct infection by SARS-CoV-2, and that the virus may induce detrimental cytopathic effects in these cells. At an MOI of 0.1, whereby approximately 1 viral particle was introduced per 10 cells, hiPSC-CMs experienced cytopathic effect after 72 hours of infection. The virus presumably enters the cell through the ACE2 receptor, which has previously been shown to be produced in hiPSC-CMs (Churko et al., 2018). Immunostaining of two SARS-CoV-2 components, the unique dsRNA intermediate and the “spike” capsid protein responsible for viral entry and virion particle assembly, demonstrates that this novel coronavirus can enter hiPSC-CMs to unleash its RNA cargo and hijack host hiPSC-CM translational machinery to produce new viral components. Notably, both the dsRNA and “spike” stains localized to perinuclear regions, indicative of the formation of hallmark viroplasm structures within hiPSC-CMs. The viroplasm is an inclusion body within the cytoplasm that recruits host cellular components such as ribosomes, chaperones, and mitochondria to establish a de facto viral replication factory within the cell (Netherton and Wileman, 2011). Not only has this perinuclear viral component accumulation been reported in other *in vitro* coronavirus infection studies (Hagemeijer et al., 2012), but it parallels what was seen in hiPSC-CMs infected with coxsackievirus B3 (Sharma et al., 2014). Presumably, this increased viral load within hiPSC-CMs activated downstream apoptotic events. This was indicated by alterations in cellular morphologies and increased production of cleaved caspase-3.

In parallel with these stains showing viral accumulation, there was a functional alteration in hiPSC-CM contractility where the cells ceased beating. The cause of this dramatic functional change in hiPSC-CM beating must be further investigated, but perhaps cytopathic effect and associated hiPSC-CM apoptosis significantly disrupted cell-cell gap junctions and therefore impaired proper excitation-contraction coupling within and between cardiomyocytes. Indeed, significant alterations in calcium handling and contractility were similarly observed in CVB3-infected hiPSC-CMs (Sharma et al., 2014). Additional work remains to be done to further elucidate the cardiomyocyte-specific effects of SARS-CoV-2, in particular impacts on cell function and gene expression.

The cellular immaturity of hiPSC-derivative cell types must be considered in this model. It is known that hiPSC-CMs are electrophysiologically, structurally, and genetically immature in comparison to their adult counterparts (Bedada et al., 2016), although biomechanical efforts to mature these cells towards an adult phenotype have recently gained traction (Ronaldson-Bouchard et al., 2018). However, even in these immature cells, the current study showed overt virus-induced cardiotoxicity and physiological effects at low SARS-CoV-2 MOI. Since expression of *ACE2* increases as hiPSC-CMs mature in *in vitro* culture (Churko et al., 2018), one could hypothesize that after maturation, older cardiomyocytes would be even more susceptible to SARS-CoV-2 infection. Significant additional data is needed to test these possibilities.

The results presented here are in an isolated *in vitro* system, devoid of any immune system components that are thought to be critical to the many aspects of the COVID-19 viral response *in vivo* (Shi et al., 2020b). The resulting myocarditis, by definition, is an inflammation of the myocardium caused by immune cell infiltration, which may or may not be the result of direct cardiomyocyte viral infection. However, as shown here, human iPSC-derived cardiomyocytes are clearly susceptible to SARS-CoV-2 infection even in the absence of an immune response, suggesting the possibility of direct infection of the heart playing a role in the clinical syndrome. Indeed, a very recent study using biopsied heart tissue shows the SARS-CoV-2 virus in the myocardium of a COVID-19 patient (Tavazzi et al., 2020). While this needs to be replicated, it raises the question of whether the SARS-CoV-2 enters the heart via a viremic phase or through macrophages clearing the virus from the lungs. Nevertheless, these studies suggest that cardiomyocyte infection may contribute to the variety of cardiac clinical sequelae presented by patients with COVID-19, such as arrhythmias, elevated cardiac troponin biomarkers, or heart failure.

We predict that future studies will be able to use hiPSC-CMs as a cardiac-specific antiviral drug screening platform, in similar fashion to our previous studies with coxsackievirus B3 on hiPSC-CMs (Sharma et al., 2014). A variety of antiviral approaches, ranging from repurposed small molecules such as nucleoside analogues or viral polymerase inhibitors to novel antibodies and antisense oligonucleotides, have been proposed for COVID-19 and are currently being tested *in vitro* and in clinical trials (Harrison, 2020). HiPSC-CMs could now serve as a cardiac-specific auxiliary cell type for *in vitro* pre-clinical efficacy studies for any drug aiming to stymie SARS-CoV-2 proliferation. In parallel, drug-induced arrhythmias and QT-interval prolongation can also be examined for antiviral compounds in pre-clinical development, given that some existing COVID-19 drug treatments exhibit these off-target cardiotoxicities (Roden et al., 2020). Due to the enormity of the current COVID-19 pandemic, and the increasing evidence for associated cardiac symptoms, innovative approaches using technologies such as the hiPSC-CMs presented here will critical to further understand and counteract SARS-CoV-2 infection.

## METHODS

### HiPSC Derivation

The appropriate institutional review board (IRB) and stem cell research oversight committee (SCRO) were consulted at UCLA and Cedars-Sinai Medical Center. The 02iCTR hiPSC line was derived from human peripheral blood mononuclear cells and has been published previously (Laperle et al., 2020). This cell line was obtained from the Cedars-Sinai Medical Center iPSC core facility and was derived under their IRB-SCRO protocol “Pro00032834: iPSC Core Repository and Stem Cell Program”.

### HiPSC-CM Differentiation

The iPSC differentiation work in the Svendsen Lab is carried out under “Pro00021505: Svendsen Stem Cell Program”, authorized by the Cedars-Sinai Medical Center IRB. The hiPSC-CMs were generated from hiPSCs using a small-molecule mediated differentiation approach that modulates Wnt signaling (Sharma et al., 2015). Briefly, this approach uses the CHIR99021 GSK3β inhibitor to initiate mesoderm specification, followed by Wnt-C59 Wnt inhibitor to initiate cardiac specification. Cells began beating at approximately day 7 post-differentiation. Cardiomyocytes were metabolically selected from other differentiated cells by using glucose deprivation as previously described (Sharma et al., 2015). After selection, hiPSC-CMs were replated as a monolayer into 96-well plate format at 100,000 cells per well in hiPSC-CM culture medium, RPMI 1640 + B27 supplement with insulin.

### SARS-CoV-2 infection of hiPSC-CMs

SARS-CoV-2, isolate USA-WA1/2020, was obtained from the Biodefense and Emerging Infections (BEI) Resources of the National Institute of Allergy and Infectious Diseases (NIAID). Importantly, all studies involving SARS-CoV-2 infection of hiPSC-CMs were conducted within a Biosafety Level 3 facility at UCLA. SARS-CoV-2 was passaged once in Vero-E6 cells and viral stocks were aliquoted and stored at −80°C. Virus titer was measured in Vero-E6 cells by TCID_50_ assay. Vero-E6 cells were cultured in DMEM growth media containing 10% fetal bovine serum, 2 mM L-glutamine, penicillin (100 units/ml), streptomycin (100 units/ml), and 10 mM HEPES. Cells were incubated at 37°C with 5% CO_2_.

For hiPSC-CM infection, viral inoculum (MOI of 0.01) was prepared using serum-free media. Culture media from each well containing hiPSC-CMs was removed and replaced with 250 µL of prepared inoculum. For mock infection, serum-free media (250 µL/well) alone was added. The inoculated plates were incubated for 1 hour at 37°C with 5% CO_2_. The inoculum was spread by gently tilting the plate sideways at every 15 minutes. At the end of incubation, the inoculum was replaced with fresh hiPSC-CM culture medium. Cells remained at 37°C with 5% CO_2_ for 72 hours before analysis.

### Imaging and Immunofluorescence

After 72 hours of SARS-CoV-2 infection (or mock), live cell images and videos were obtained by bright field microscope (Leica DMIL LED). For immunofluorescence, separate wells of hiPSC-CMs were fixed with 4% paraformaldehyde in phosphate-buffered saline (PBS) for 20 minutes. The fixed samples were then permeabilized and blocked for 1 hour in a “blocking solution” containing PBS with 2% bovine serum albumin, 5% donkey serum, 5% goat serum, and 0.3% Triton X-100. Primary antibodies were diluted in the blocking solution and added to samples overnight at 4°C. The following antibodies and dilutions were used: α-actinin (1:100, Sigma-Aldrich Cat# A7811, RRID: AB_476766); cardiac troponin T (cTnT, 1:100, Abcam Cat# ab45932, RRID: AB_956386); SARS-CoV-2 spike (S) protein (1:100, BEI Resources NR-616 Monoclonal Anti-SARS-CoV S Protein (Similar to 240C) SARS coronavirus); SARS-CoV-2 double stranded RNA (1:100, J2 clone; Absolute Antibody Inc.); cleaved caspase-3 (1:200, Cell Signaling Technology Cat# 9661, RRID: AB_2341188). Samples were then rinsed 5 times for 2 minutes each with PBS containing 0.3% Triton X-100, followed by incubation with fluorescent-conjugated secondary antibodies diluted 1:1000 in blocking buffer for 2 hours at room temperature. Antibodies were donkey anti-rabbit 488 (Thermo Fisher Scientific Cat# R37118, RRID: AB_2556546), goat anti-mouse 555 (Thermo Fisher Scientific Cat# A32727, RRID: AB_2633276), and donkey anti-mouse 594 (Thermo Fisher Scientific Cat# R37115, RRID: AB_2556543). Samples were then rinsed 5 times for 2 minutes each with PBS containing 0.3% Triton X-100, followed by DAPI diluted in PBS at 1:5000 for 10 minutes. Immunofluorescence images were quantified using ImageJ software. DAPI was used to count total cell numbers in order to obtain a percentage of cells positive for dsRNA, spike protein, or cleaved caspase-3.

### Beat rate analysis

Thirty-second video clips were taken of mock and infected hiPSC-CMs at 72 hours after viral infection. A beats-per-minute measurement was obtained by manually counting individual beats in a 30-second video and multiplying by 2 to extrapolate to 1 minute.

### Statistics

For statistical analyses, the Student’s t test was used for comparison between two datasets. All error bars refer to standard deviation. A p value of <0.05 is considered statistically significant.

## AUTHOR CONTRIBUTIONS

A.S., V.A., and C.N.S. designed analyses, analyzed data, and drafted the manuscript. A.S. and G.G. conducted experiments and acquired data. A.S., V.A., and C.N.S. analyzed data. All authors contributed to the final manuscript.

## ACKNOWLEDGEMENTS AND FUNDING

We acknowledge that this review represents an area of study that is dramatically growing in relevance, and thus we apologize to any authors whose work we were not able to include here. We thank Dr. Soshana Svendsen for editing the manuscript. Research from the Svendsen laboratory has been supported by the National Institutes of Health (5UG3NS105703) and the Cedars-Sinai Board of Governor’s Regenerative Medicine Institute. Arun Sharma is supported by an institutional training grant (T32 HL116273). The following reagent was obtained through BEI Resources, NIAID, NIH: Monoclonal Anti-SARS-CoV S Protein (Similar to 240C), NR-616.

